# Identification of markers for the isolation of neuron-specific extracellular vesicles

**DOI:** 10.1101/2024.04.03.587267

**Authors:** Dmitry Ter-Ovanesyan, Sara Whiteman, Tal Gilboa, Emma JK Kowal, Wendy Trieu, Siddharth Iyer, Bogdan Budnik, Clarissa May Babila, Graham Heimberg, Michael W Burgess, Hasmik Keshishian, Steven A Carr, Aviv Regev, George M Church, David R Walt

**Affiliations:** Wyss Institute for Biologically Inspired Engineering, Boston, MA, USA; Department of Biology, Massachusetts Institute of Technology, Cambridge, MA, USA; Department of Biological Engineering, Massachusetts Institute of Technology, Cambridge, MA, USA; Genentech, South San Francisco, CA, USA; Broad Institute of MIT and Harvard, Cambridge, MA, USA; Harvard Medical School, Boston, MA, USA; Department of Pathology, Brigham and Women’s Hospital, Boston, MA, USA

## Abstract

Extracellular vesicles (EVs) are released by all cells and contain RNA and protein from their cell of origin. EVs in biofluids could be used as diagnostic biomarkers to non-invasively report the state of inaccessible cells, such as neurons in the brain. As biofluids such as cerebrospinal fluid (CSF) and plasma contain EVs originating from many different cells, isolating cell type-specific EVs and measuring their cargo could help determine the state of specific cell types. Here, we demonstrate an approach aiming to immuno-isolate EVs from neurons based on neuron-derived protein surface markers. We first developed a framework to select transmembrane proteins suitable as neuron-specific EV markers based on gene expression and EV proteomics data. Leveraging a novel, high-purity EV isolation method we developed, we further cataloged the proteins present on EVs in human CSF and plasma. Using ultrasensitive immunoassays against several of the predicted neuron-specific proteins, we confirmed one marker, NRXN3 as present on EVs in CSF and plasma by size exclusion chromatography (SEC) and density gradient centrifugation (DGC). Finally, we developed efficient EV immuno-isolation methods and applied them to isolate NRXN3^+^ EVs. Our study provides a general methodology for the isolation of cell-type specific EVs and paves the way for the use of neuron-derived EVs to study and diagnose neurological disease.

## Introduction

Extracellular vesicles (EVs) are released by all cells into biological fluids. Since EVs contain RNA transcripts and proteins from their donor cells, they represent a rich source of biomarkers that promises to provide molecular information from inaccessible tissues and organs. However, while the total population of EVs can be isolated and profiled from plasma or other biological fluids in bulk, such profiles do not distinguish cell type of origin of these cargoes (1). Measuring the cargo of EVs from a specific cell type, on the other hand, could provide a non-invasive “snapshot” reflecting the state of those cells (2). This could in principle be done by immuno-isolation using an antibody against a transmembrane protein marker that is present on EVs and specific to that cell type.

Isolating cell-type specific EVs derived from neurons or other brain cell types would be particularly useful, given the limited accessibility of the brain to biopsy, and could shed light on diverse neurological conditions, from inflammatory (3) to neurodegenerative (2, 4) and neuropsychiatric disease (5). Multiple studies have reported the isolation and analysis of neuron-derived EVs in the context of neurological disease, mostly by immuno-isolation with antibodies against L1CAM (2, 6). However, we previously showed that L1CAM is present in plasma and CSF as a free protein, and not associated with EVs (7), highlighting the need for other candidate markers. Although other markers, such as NCAM1 (8, 9), GluR2 (10, 11), and, more recently, ATP1A3 (12) and MAP1B (13) have also been suggested, the presence of these markers on EVs has not been tested with isolation methods that can discriminate EV proteins from contaminating free proteins (14).

Here, we developed a general framework for selecting cell type-specific EV markers and applied this framework to the isolation of neuron-specific EVs. We first assessed all human proteins for whether they have a transmembrane domain, are neuron-specific, and are present on EVs in biofluids. To facilitate this query, we developed a novel high-purity EV isolation method, which enabled us to create reference proteomics profiles for human CSF and plasma EVs. We then identified which of the transmembrane proteins present on EVs are neuron-specific within the brain and brain-specific relative to other organs, based on reference gene expression atlases at the bulk and single cell levels, yielding a list of candidate neuron-specific EV markers. We tested several of these candidates and validated one such marker, NRXN3, as present on EVs in human CSF and plasma. Finally, we developed highly efficient and specific EV immuno-isolation methods, which we used to isolate NRXN3^+^ EVs from both human CSF and plasma. Our study demonstrates the use of a candidate marker for the isolation of cell type-specific EVs from neurons and presents a systematic approach that could readily be applied to EVs from other cell types.

## Results

### General framework for cell type-specific EV marker selection

To isolate neuron EVs from human biofluids (**Fig. 1A**), we first set out to develop a general framework for the identification of cell type-specific EV markers. We reasoned that each candidate marker should satisfy three criteria: (**1**) be present on EVs in the biofluid of interest; (**2**) be a transmembrane protein, a pre-requisite for EV immuno-isolation; and (**3**) be specifically expressed in the cell type of interest relative to other cell types. To help test for these criteria, we developed a method to isolate high-purity EVs and profile them by proteomics (to assess (1)), as well as a computational pipeline that relies on reference bulk and single cell RNA-seq (scRNA-Seq) to identify tissue- and cell-type specific genes that encode transmembrane proteins (to address (**2**) and (**3**)), as we present below. We developed and applied both methods with the goal of isolating neuron-specific EVs.

**Figure 1:**
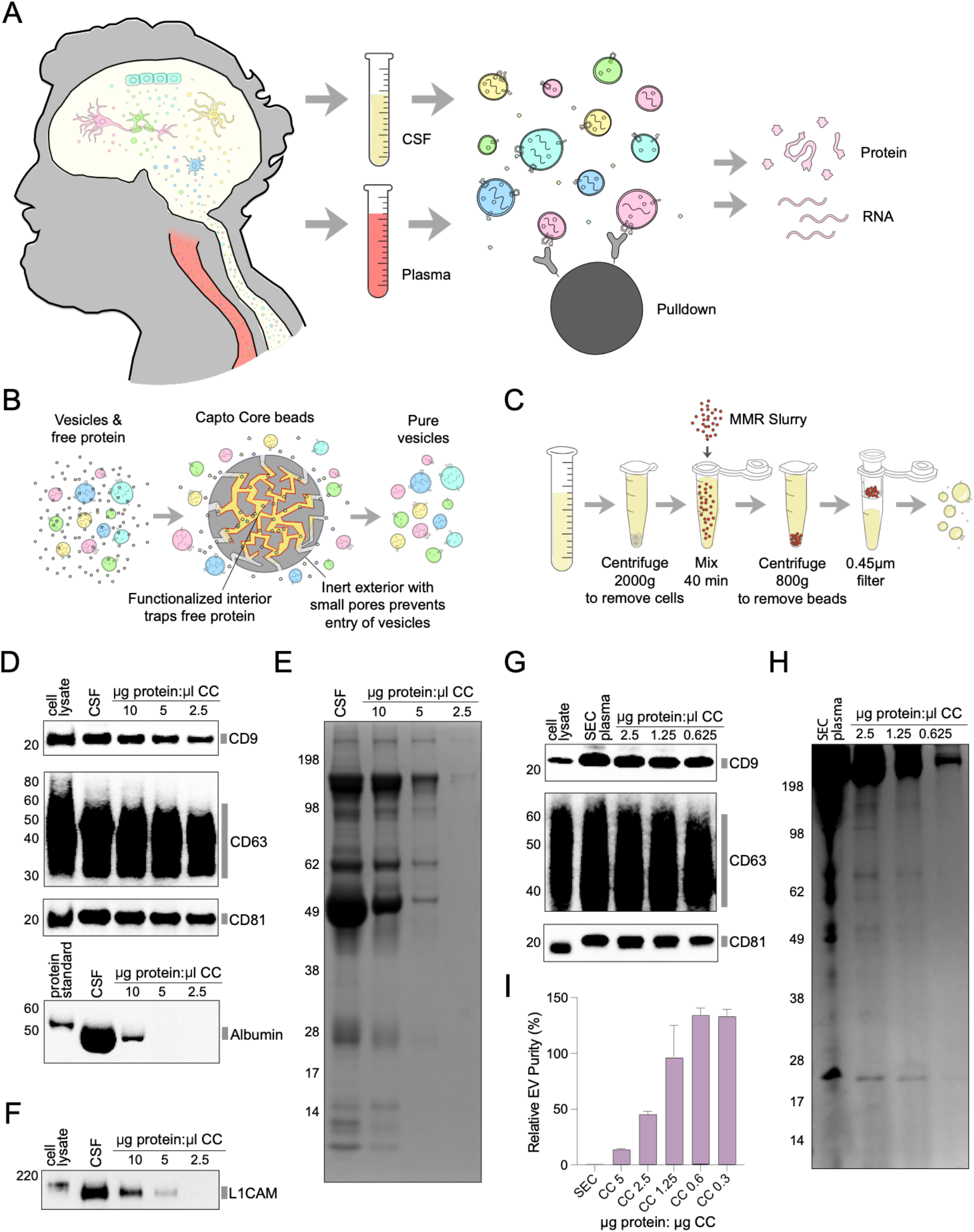
Development of MMR Slurry for high purity isolation of extracellular vesicles from human CSF and plasma. **A.** Overview of approach. Immuno-isolation of neuron-specific EVs from human CSF or plasma for interrogation of neuron-derived RNA or protein cargo. **B.** Principle underlying MMR Slurry technique. Capto Core 700 beads have pores allowing biomolecules less than 700 kDa to enter, and once free proteins enter, they stay in the beads, which can be removed to leave pure EVs. **C.** EV isolation workflow with MMR Slurry. Samples are mixed with Capto Core beads and then beads are pelleted and discarded. **D.** Western blot of CD9, CD63, CD81, and albumin of human CSF after MMR Slurry purification with increasing amounts of Capto Core beads shows a strong enrichment of tetraspanins relative to albumin. **E.** Total protein stain of CSF after MMR Slurry purification with increasing amounts of Capto Core beads. **F.** Western blot of L1CAM in CSF after MMR Slurry purification with increasing amounts of Capto Core beads. **G.** Western blot of CD9, CD63, CD81, and albumin after SEC of plasma followed by MMR Slurry purification with increasing amounts of Capto Core beads shows a strong enrichment of tetraspanins relative to albumin. **H.** Total protein stain after size exclusion chromatography of plasma, followed by MMR Slurry purification with increasing amounts of Capto Core beads. Ratio of 50% Capto Core (CC) bead slurry to protein is indicated in μL CC Slurry/μg protein in sample. For D-H, equal fraction each sample was loaded onto each gel. **I.** Simoa quantification of EV purity comparing MMR Slurry after SEC or SEC from plasma. Levels of CD9, CD63, and CD81, and albumin were measured between conditions (based on two replicates for each condition and Simoa measurements made in duplicate for each replicate). Ratios of CD9, CD63, and CD81 were averaged to calculate relative EV yield and divided by relative ratios of albumin to calculate EV purity. EV purity of SEC is set to 1. Two separate replicates of all samples were used and Simoa measurements were taken in duplicate for each sample. Error bars indicate SD.

### Development of a high-purity EV isolation method for proteomics

We first aimed to use proteomic profiling to catalog all proteins present on EVs in human CSF and plasma. Because free proteins such as albumin are many orders of magnitude more abundant than EV proteins in biofluids, their presence in EV preparations limits the ability of mass spectrometry to detect EV proteins (15). Although we have previously optimized size exclusion chromatography (SEC) to reduce free proteins such as albumin by several orders of magnitude, the EV fractions in SEC still carry substantial levels of albumin (16).

We thus developed new methods for purifying EVs from human biofluids with very high purity to maximize the depth of protein coverage by mass spectrometry. We chose a mixed mode chromatography (MMC) resin (Capto Core 700, **Methods**), comprised of beads with an inert outer shell and pores that exclude molecules larger than 700 kDa. The MMC beads’ core contains octylamine ligands that are both hydrophobic and positively charged, efficiently trapping proteins that enter the beads. This resin was originally developed for viral purification in chromatography columns (17), and we have previously shown that it can be added to the bottom layer of SEC columns (18). Inspired by the use of MMC resin for viral purification “in slurry” without the columns for non-enveloped, infectious virus purified from cells (19), we tested MMC resin for EV purification.

We developed a straightforward method, Mixed Mode Resin (MMR) Slurry, where MMC resin is mixed with biofluids to purify EVs away from free proteins (**Fig. 1B,C**). MMR Slurry is based on the insight that since MMC beads bind and trap free proteins, the beads could be separated after incubation with the biofluid, leaving pure EVs in solution (**Fig. 1B,C**). To optimize MMR Slurry, we measured the levels of the tetraspanins CD9, CD63, and CD81 (as a proxy for EV yield) *vs*. albumin (for free protein contamination) (**Fig. 1D**), finding that the ratio of resin to total protein was a crucial parameter. Applying MMR Slurry to CSF, we found that by increasing the ratio of resin volume to total protein, we could reproducibly “tune” EV purity. In this way, we were able to eliminate albumin (as measured by Western blot), while retaining a large fraction of the EVs (**Fig. 1D,E**). In agreement with our previous results that L1CAM is not an EV marker in CSF (7), L1CAM also completely disappeared by Western blot upon increasing MMR amounts (**Fig. 1F**). For plasma, which has two orders of magnitude more free protein than CSF, we first purified plasma using SEC and then applied MMR Slurry to 1 mL of plasma, depleting albumin to almost undetectable levels (**Fig. 1G, H**).

To quantify the increase in EV purity enabled by MMR Slurry, we used previously developed Single Molecule Array (Simoa) immunoassays against CD9, CD63, CD81, and albumin (16, 18). Simoa measures single protein molecules in individual microwells and, thereby, turns ELISA into a highly sensitive, digital readout. When averaging the ratios of tetraspanins between conditions and comparing these to albumin levels, MMR Slurry improved EVs purity more than 100-fold compared to SEC (**Fig. 1I, Fig. S1**). Thus, MMR Slurry could be used to isolate EVs with high purity from CSF as a simple one-step purification or from plasma as a two-step protocol after SEC.

### Mass spectrometry of EVs from human plasma and CSF using MMR Slurry

We applied MMR Slurry to profile the EV proteome for human pooled CSF and plasma. Using the one-step MMR Slurry method for CSF or the two-step SEC and MMR Slurry method for plasma (**Fig. 2A**), we performed mass spectrometry on EVs isolated from 1 mL of biofluid. To remove lipoproteins, we also found that MMR Slurry can be performed after reducing ApoB100 levels by replacing SEC with dual mode chromatography (DMC), which combines SEC with cation exchange resin (20). Combining data from several mass spectrometry runs across pooled CSF or plasma isolated by MMR Slurry, we generated reference EV proteomes of 2,104 proteins for CSF and 1,862 proteins for plasma (**Fig. 2B, Table S1**).

**Figure 2:**
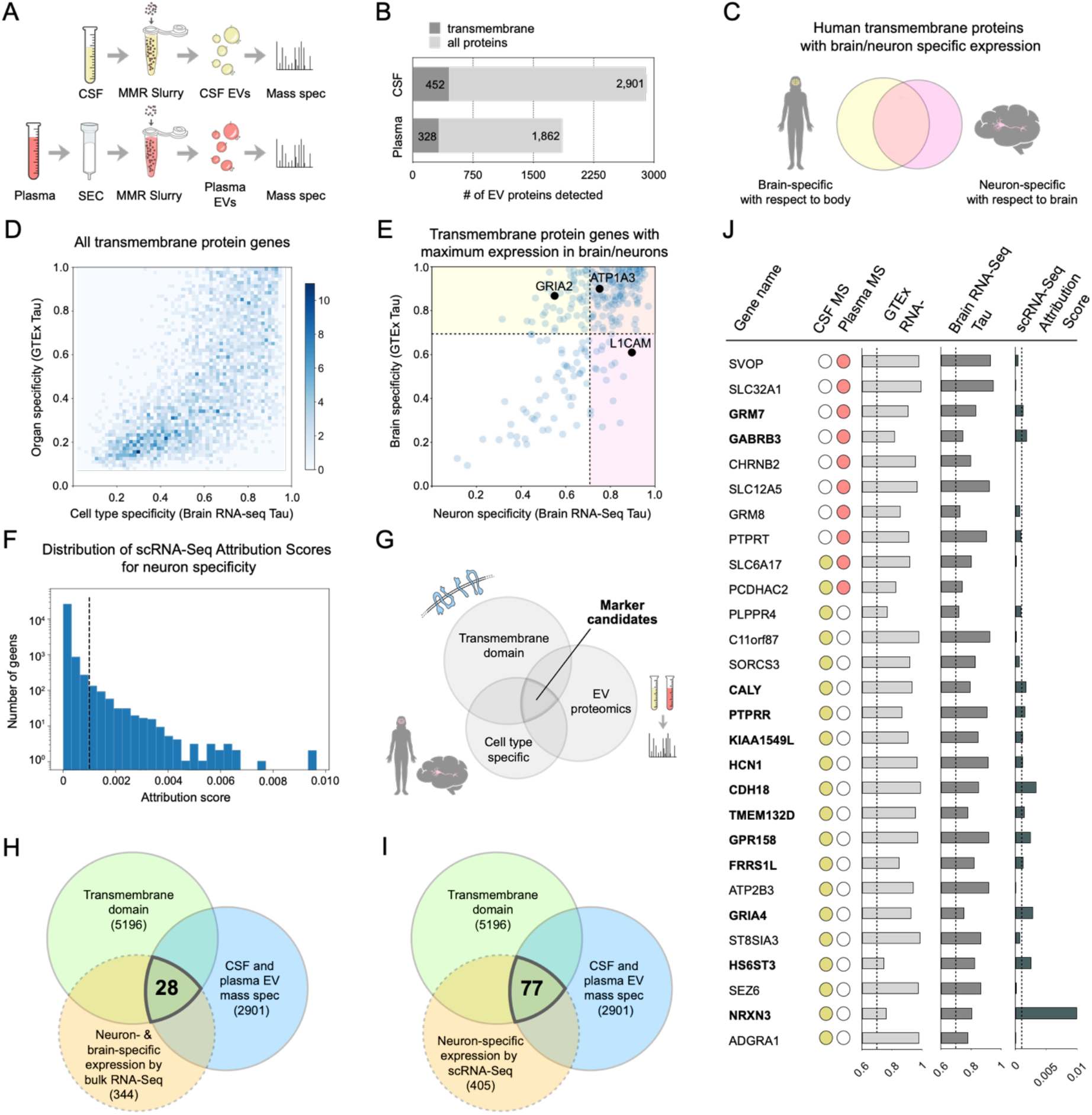
Identifying candidate markers for isolation of neuron-specific extracellular vesicles using gene expression and EV proteomics data. **A.** Schematic of EV proteomics using MMR Slurry for CSF and SEC followed by MMR Slurry in plasma. **B.** Number of distinct total proteins and transmembrane proteins detected in pooled human CSF and plasma. **C.** To find candidate markers that are cell-type specific using bulk RNA-Seq datasets, genes that specific to the brain relative to other organs are overlapped with genes that are specific to neurons relative to other cells of the brain. **D.** Tau specificity scores for brain cell type (x axis) and organ (y axis) for all transmembrane protein genes. **E.** Tau specificity scores for neurons (x axis) and brain (y axis) for transmembrane protein genes that are maximally expressed in neurons relative to other cell types of the brain and the brain relative to other organs. Tau threshold of 0.7 or above is indicated. Previously used neuron EV markers L1CAM, GRIA2, ATP1A3 are highlighted. **F.** Distribution of Attribution Scores for neuron-specific gene expression across single cell RNA-Seq studies using SCimilarity for all genes. **G.** Overview of pipeline for identification of cell type-specific EV markers: assessing transmembrane proteins for neuron-specific gene expression (based on either bulk or single cell RNA-Seq) and presence on EVs (from proteomics data). **H.** Using a Tau cutoff of 0.7 for neuron and brain-specific expression based on bulk RNA-Seq data together with transmembrane proteins found on EVs in MMR Slurry proteomics data yields 27 candidate neurons-specific EV markers. **I.** Using an Attribution Score cutoff of 0.001 for neuron-specific expression based on single cell RNA-Seq data together with transmembrane proteins found on EVs in MMR Slurry proteomics data yields 34 candidate neurons-specific EV markers. **J.** List of 27 candidate neuron-specific EV markers with their Tau values for specificity of expression in neurons relative to other cell types of the brain and the brain relative to other organs in the body, SCimilarity scRNA-Seq Attribution Score, as well as presence in CSF and/or plasma EV MMR mass spectrometry data. The 27 candidate markers listed all made the specificity score cutoff based on bulk RNA-Seq. The subset of these that also made the specificity score cutoff based on scRNA-Seq is indicated in bold.

### Gene expression-based selection of neuron-specific EV marker candidates

To determine which of the EV proteins detected in CSF and plasma could be candidates for cell type-specific EV isolation, we analyzed all human proteins for both annotated transmembrane domains (21) in UniProt (5,196 of 20,375 UniProt proteins in UniProt, **Table S2**), and gene expression in the desired cell type. We selected neuron-specific genes based on RNA-Seq of the major cell types isolated from human brain tissue via immuno-panning (22)) and tissue (based on GTEx RNA-Seq (23)) (**Fig. 2C**), using the gene expression specificity index, Tau (24), as a robust measure (25) of each gene’s specificity (**Fig. 2D**). This approach identified 165 transmembrane proteins that are expressed at higher levels in neurons *vs*. other brain cells and in brain *vs*. other organs, while meeting our specificity thresholds (Tau > 0.7 for each) (**Fig. 2E, Fig. S2, Table S3**), as an initial set of candidates.

In parallel, we queried scRNA-Seq profiles of >22 million cells from 399 studies using SCimilarity (26), a deep learning model that allows us to rank cell-type specificity across scRNA-Seq datasets, which are otherwise challenging to compare (27, 28). SCimilarity assigned an attribution score to each gene indicating how neuron-specific its expression is. Based on the distribution of SCimilarity’s attribution scores across all genes (**Fig. 2F**), we chose a cutoff of 0.001, yielding 405 neuron-specific genes, of which 179 encode proteins with annotated transmembrane domains (**Table S4**). Of these 179, 113 genes are shared with the 165 identified from the bulk RNA-Seq datasets (**Table S5**).

Finally, we selected candidate neuron-specific EV markers as those genes that encode proteins with annotated transmembrane domains, have neuron-specific expression, and are present on EVs in CSF or plasma by proteomics (**Fig. 2G-J**). We identified 28 candidates when using bulk RNA-Seq for specificity in brain and neurons (**Fig. 2H**), 77 candidates when using scRNA-Seq (**Fig. 2I**), and 13 in both (**Fig. 2J**). In addition to our proteomics data, we also analyzed other existing mass spectrometry datasets (29-39). We also applied the same modular pipeline to identify candidate cell type-specific EV markers for other cell types in the brain (**Table S6**).

### Validation of neuron-specific EV markers

Next, we set out to confirm that the candidate markers are present on EVs, to avoid immuno-isolation of soluble protein isoforms not associated with EVs, as we previously found for L1CAM (7). Based on antibody availability, we selected three markers for testing: SLC12A5, SLC32A1, and NRXN3. We developed Simoa immunoassays for each protein, compared all pairs of commercially available antibodies to each other, and using the best antibody pair, validated the performance of each assay in CSF and plasma (**Fig. S3-S5, Table S7**). To test our assays, we also used a positive control system based on an established human induced pluripotent stem cell (iPSC) line that rapidly and efficiently differentiates into neurons upon doxycycline-inducible expression of the transcription factors (TFs) Neurogenin1/2 (NGN) (40) (**Fig. S6**). We performed SEC on CSF, plasma, or EVs enriched by ultracentrifugation (41) from the culture media of induced Neurogenin1/2 (iNGN) cells and measured the levels of the three markers in each of the SEC fractions with Simoa (7, 16). We detected NRXN3 in the early SEC fractions of neuron EVs, CSF, and plasma (**Fig. 3A,B, Fig. S5**). SLC12A5 was not detected in any of the early SEC fractions (**Fig. S3**), and SLC32A1 peaked only in the early SEC fractions of the iNGN neuron EVs but not CSF or plasma (**Fig. S4**)

**Figure 3:**
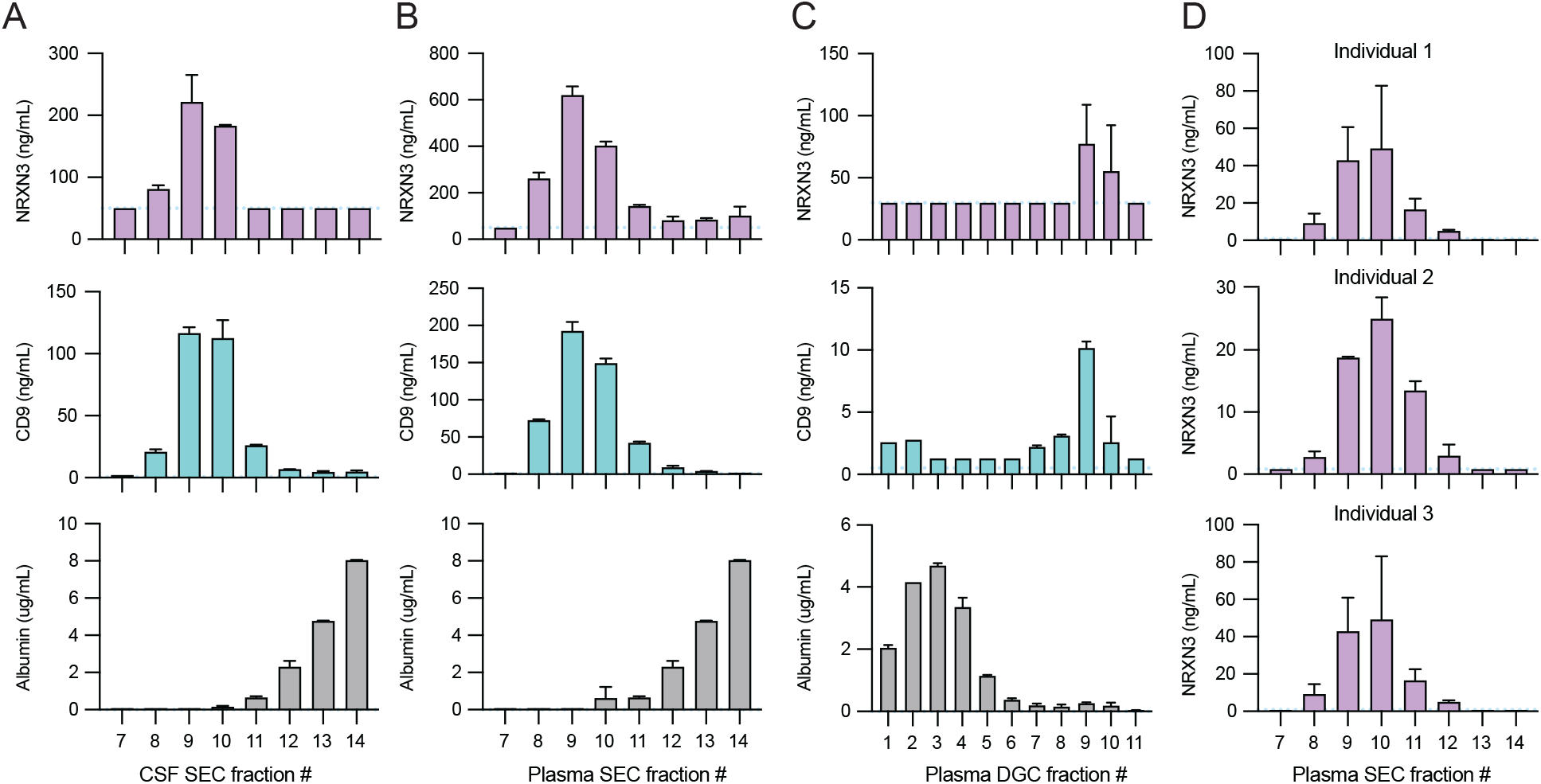
Validation of NRXN3 as neuron-specific marker present on EVs and immuno-isolation of NRXN3^+^ EVs. **A.** Simoa measurement in SEC fractions of pooled CSF of NRXN3 (top), CD9 (middle) and albumin (bottom). **B.** Simoa measurement in SEC fractions of pooled plasma of NRXN3 (top), CD9 (middle) and albumin (bottom). **C.** Simoa measurement in DGC fractions of pooled plasma of NRXN3 (top), CD9 (middle) and albumin (bottom). **D.** Simoa measurement of NRXN3 in SEC fractions of plasma from three different individuals. For A-D, Simoa measurements were taken in duplicate for each sample. Error bars indicate SD.

We further validated NRXN3 on CSF and plasma EVs based on different isolation and/or detection methods, as well as individual, rather than pooled, samples. First, we performed density gradient centrifugation (DGC), and measured each fraction by Simoa. NRXN3 was present in the same fractions as CD9, but not albumin, in two different batches of pooled plasma (**Fig. 3C**). In CSF, the DGC results varied between two batches with NRXN3 present in the early fractions in one batch but later fractions in the other (**Fig. S7**). Second, we detected NRXN3 by Western blotting after enriching for EVs using MMR Slurry (**Fig. S8**). Finally, we performed SEC on three individual CSF and plasma samples and detected NRXN3 by Simoa in the early SEC fractions of each of the samples (**Fig. 3D, Fig. S9**). Thus, we confirmed that NRXN3 is present on EVs from iNGN neurons, CSF, and plasma, which is particularly important since NRXN3 has also been reported to have soluble isoforms (42, 43). NRXN3 is highly specific to neurons in both bulk and scRNA-Seq datasets (Fig. 2J), and we further confirmed it is highly expressed across different subtypes of neurons in human brain scRNA-Seq dataset (44) (**Fig. S10**).

### Optimization of EV immuno-isolation with EVs enriched from cell culture media

Since EVs from neurons containing a marker such as NRXN3 are likely to be a small fraction of all EVs in a biofluid such as plasma, we next developed and validated a highly efficient and specific protocol for immuno-isolation of EV subsets. We initially optimized a protocol in the human K562 cell line using the widely expressed tetraspanin CD81. We purified EVs from the conditioned media of K562 cells using differential ultracentrifugation (41), performed immuno-isolation, and evaluated the efficiency via Western blotting of CD81 in the pulldown (PD) *vs*. flow-through (FT) fraction (14) (**Fig. 4A**). To ensure that our immuno-isolation is specific for the target protein, we also measured CD81 when EVs were immuno-isolated using the same procedure but with a non-specific antibody. We systematically optimized several parameters, including antibody conjugation strategy, bead and antibody quantity and ratio (**Fig S11**). Our final protocol led to immuno-isolation with high efficiency and specificity (**Fig. 4B)**. To more closely model our final goal of isolating neuron EVs, we also used iNGN cell EVs. To characterize these EVs, we isolated them from the cell culture media of iNGN cells and characterized them by mass spectrometry together with EVs from the parental iPSC line (**Table S8**). To model cell-type specific EV capture from biofluids, we further optimized the protocol on neuron EVs (**Fig. S12, S13**) and then mixed neuron EVs with iPSC EVs and confirmed that our immuno-isolation is specific for neuron but not iPS EVs, even with a low (1:10) proportion of neuron EVs relative to non-neuron EVs (**Fig. S14**).

**Figure 4:**
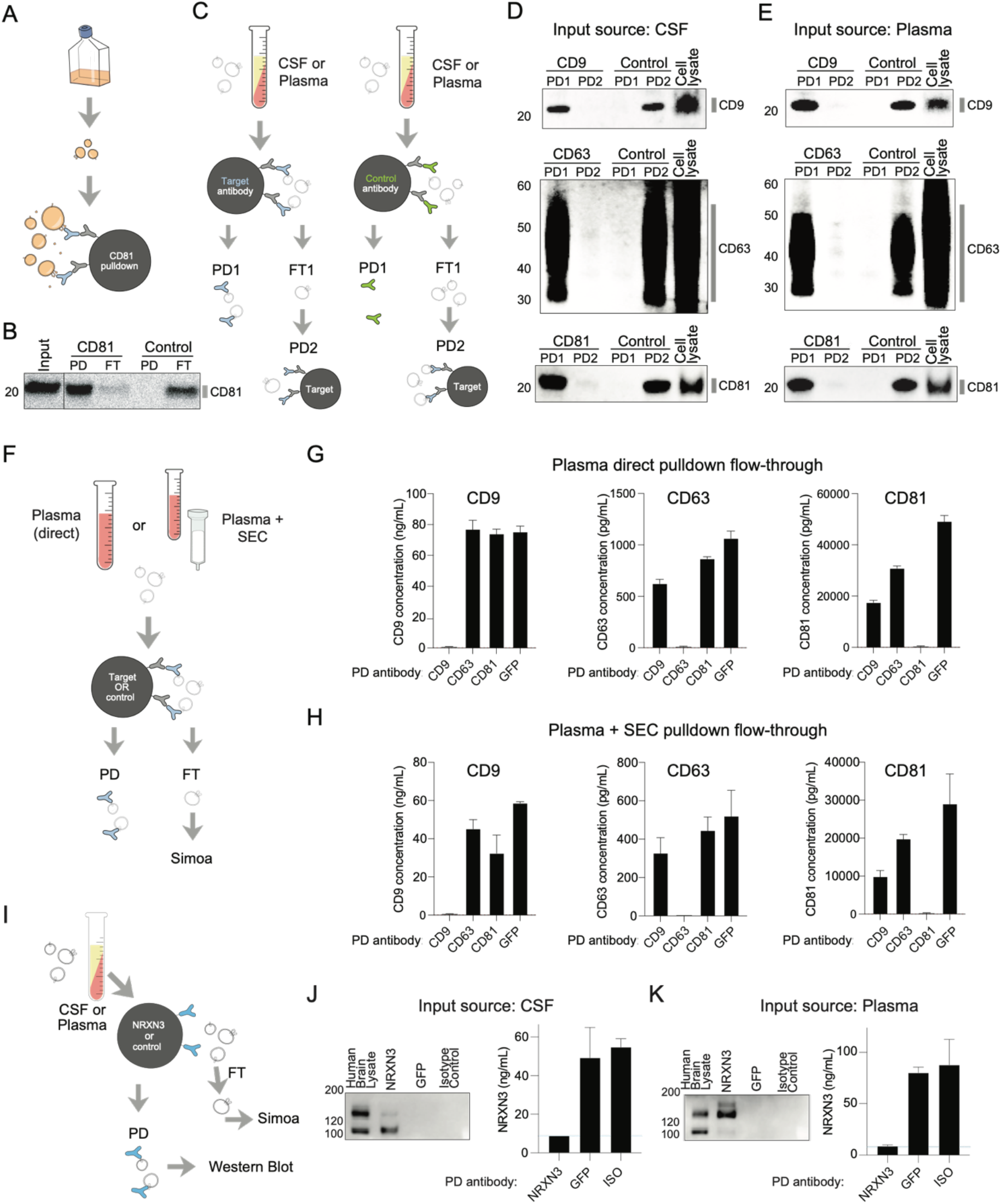
Development and validation of EV immuno-isolation method and application to NRXN3. **A.** Schematic of immuno-isolation of EVs enriched by ultracentrifugation from the cell culture media of K562 cells. **B.** Western blotting of CD81 after immuno-isolation of EVs. Equal amounts of EVs obtained by differential ultracentrifugation of K562 cell culture media were loaded as input or immuno-isolated using either an anti-CD81 antibody or a control non-specific antibody. PD = pulldown, FT = flow-through **C.** Schematic of EV immuno-isolation experiment from human plasma or CSF for western blotting analysis. EVs are immuno-isolated from biofluid without initial EV purification. Immuno-isolation was performed using a target antibody or control antibody (PD1 = pulldown 1) and the flow-through is subjected to a second immuno-isolation with the target antibody (PD2 = pulldown. 2) **D.** Western blots of CD9 (top), CD63 (middle), or CD81 (bottom) after immuno-isolation from human CSF. In each case, the first pulldown (PD1) was performed with antibodies against the target protein or a control antibody, and the second pulldown (PD2) was performed with antibodies against the target protein using the flow-through of PD1. **E.** Western blots of CD9 (top), CD63 (middle), or CD81 (bottom) after immuno-isolation from human plasma. In each case, the first pulldown (PD1) was performed with antibodies against the target protein or a control antibody, and the second pulldown (PD2) was performed with antibodies against the target protein using the flow-through of PD1. **F.** Schematic of EV immuno-isolation experiment from human plasma or EVs enriched by SEC from human plasma for assessment by Simoa in the flow-through (FT). **G.** Simoa measurements of CD9 (left), CD63 (middle), or CD81 (right) of the flow-through of human plasma after immuno-isolation with antibodies against CD9, CD63, CD81 or GFP (non-specific control). **H.** Simoa measurements of CD9 (left), CD63 (middle), or CD81 (right) of the flow-through of EVs enriched from human plasma by SEC after immuno-isolation with antibodies against CD9, CD63, CD81 or GFP (non-specific control). For G and H, two separate replicates of all samples were used and Simoa measurements were taken in duplicate for each sample. Error bars indicate SD. **I.** Schematic of NRXN3 immuno-isolation experiment. NRXN3 in pulldown fraction (PD) was measured using Western blotting. NRXN3 in flow-through (FT) was measured using Simoa. Immuno-isolation experiment was performed using antibody against target (NRXN3) or control (GFP or isotype control) **J.** Western blotting of NRXN3 after pulldown from CSF using either NRXN3 antibody or control antibody (GFP or isotype control). Simoa measurement of NRXN3 in flow-through after pulldown from CSF using either NRXN3 antibody or control antibody (GFP or isotype control). **K.** Western blotting of NRXN3 after pulldown from plasma using either NRXN3 antibody or control antibody (GFP or isotype control). Simoa measurement of NRXN3 in flow-through after pulldown from plasma using either NRXN3 antibody or control antibody (GFP or isotype control). For J and K, Simoa measurements were taken in duplicate for each sample. Error bars indicate SD.

### Immuno-isolation of EV subsets from human CSF and plasma without prior EV purification

We next tested our optimized immuno-isolation in biofluids, by immuno-isolating EVs containing the tetraspanins CD9, CD63, and CD81 from human CSF and plasma. As in the cell culture experiments, in the first pulldown (PD1), we performed immuno-isolation using either a target or control antibody. Because the flow-through from this initial pulldown (FT1) contained protein levels that were too high to run on a protein gel, we subjected the flow-through to a second immuno-isolation with the target antibody (PD2) (**Fig. 4C**). Comparing the results of the first and second immuno-isolation (PD1 *vs*. PD2) suggested that our protocol is highly efficient and specific in both CSF (**Fig. 4D**) and plasma (**Fig. 4E**). Thus, given sufficiently specific antibodies, we can successfully perform immuno-isolation from human biofluids without first purifying EVs. To further assess our immuno-isolation, we used the Simoa tetraspanin assays to measure the depletion of the target protein in the flow-through fraction after pulldown (**Fig. 4F**). We performed our optimized immuno-isolation protocol on either total plasma (**Fig. 4G**) or EVs enriched from plasma by SEC (**Fig. 4H**) and showed that, in both cases, we achieved almost total depletion of the target protein in the flow-through fraction for each of the three tetraspanins.

### Immuno-isolation of NRXN3^+^ EVs

Finally, we immuno-isolated NRXN3^+^ EVs. Using the optimized immuno-isolation method and testing framework we developed for tetraspanins, we measured NRXN3 in both the pulldown (by Western blot) and flow-through (by Simoa) (**Fig. 4I**). We successfully and specifically immuno-isolated NRXN3 from both CSF and plasma, as indicated by enrichment in the pulldown and depletion in the flow-through compared to both an isotype control antibody and another isotype-matched, non-specific antibody against GFP (**Fig. 4J,K**). Immuno-isolation using EVs enriched from CSF or plasma by SEC was also NRXN3-specific (**Fig. S15**), further validating that NRXN3 is present on EVs. Thus, we were able to demonstrate EV immuno-isolation from both human CSF and plasma using one of our predicted neuron-specific EV candidate markers, NRXN3.

## Discussion

In this work, we provide a systematic framework for the isolation of cell type-specific EVs. This includes identification of candidate cell type-specific EV markers from gene expression and EV proteomics data, validation that a candidate marker is present on EVs, and development of optimized EV immuno-isolation methods. We apply this framework to identify neuron-specific EV markers, and after validating that one such candidate marker, NRXN3, is present on EVs in both CSF and plasma, perform NRXN3^+^ EV immuno-isolation.

Several technological advances in our work should be broadly useful for the EV field. Our computational pipeline allows one to comprehensively search for cell-type specific EV markers by analyzing pan-body RNA-Seq atlases across bulk (GTEx (23)) and single cell (Human Cell Atlas (26)) datasets. As the Human Cell Atlas (45) continues to grow (27, 28) it will be possible to further refine such searches. In this study, we treated neurons as a single “cell type,” but our parameters could be modified to find markers of neuronal subtypes. Our pipeline integrates cell type-specific transmembrane proteins with human CSF and plasma EV proteomics data we have generated using MMR Slurry, a new EV isolation method. MMR Slurry can purify EVs from clinically relevant volumes of CSF by simply incubating the resin with the biofluid, and can also increase purity ∼100 fold (**Fig. 1I**) when applied to plasma after SEC or DMC (to remove lipoproteins). As Capto Core 700 resin has recently been shown to remove dye when used in 96-well filter plates (46), we envision MMR Slurry being adaptable to a high-throughput format and well-suited for profiling EVs from clinical samples for biomarker discovery. We also optimized and validated an EV immuno-isolation protocol that works in both plasma and CSF with or without prior EV enrichment. This is crucial for the isolation of cell type-specific EVs, which may comprise a small subset of total EVs in a given biofluid.

Our study demonstrates a proof of concept for the isolation of neuron-specific EVs, but further work is necessary to validate any candidate marker, including NRXN3. As many transmembrane proteins have alternative isoforms due to alternative splicing (47) or proteolytic cleavage (48, 49), it is important to verify that a candidate EV marker is truly present on EVs. While we confirmed that NRXN3 is present on both plasma and CSF EVs by several methods, NRXN3 has many alternatively spliced isoforms, some of which are soluble (42, 43). Thus, although the Simoa assay recognized NRXN3 on EVs, it is possible that soluble forms also exist, and these could also be recognized by the antibody used for immuno-isolation. Further detailed mapping of different isoforms of NRXN3 in CSF and plasma (*e.g.*, using targeted mass spectrometry-based methods (50)) would help confirm whether a given epitope used for immuno-isolation is present exclusively on EV-bound NRXN3 or also present on soluble NRXN3.

Further studies are also required to characterize the neuron-specificity, disease-relevance, and molecular characteristics of EVs isolated using NRXN3 or other candidate markers. First, it is important to verify that any candidate marker is present at sufficiently high levels across individuals with disease status and age, and that it has the correct topology on EVs. This is important since some of our candidate markers are also present on synaptic vesicles, where they would be expected to have the opposite topology relative to EVs. Second, NRXN3^+^ EVs will need to be further analyzed further to confirm their neuronal source, for example by the enrichment for neuronal RNAs or proteins. Enzymatic treatment with proteinase or proteinase and then RNAse could help ensure that these neuron-specific cargo molecules are truly inside EVs and not non-specifically stuck to the outside (51).

Profiling the molecular cargo of neuron-specific EVs presents exciting opportunities for better understanding of brain pathology during neurodegeneration or other disease states and, especially if plasma can be used as a source, early disease detection. The resources and framework we introduce in this work represent significant steps towards this goal. Additionally, our work is broadly applicable to identifying and validating cell type-specific EV markers for other cell types and organs. We envision that this will enable novel methods to non-invasively read out cell state in health and disease.

## Acknowledgments

The authors acknowledge members of the Church, Regev and Walt labs for helpful discussions and, in particular, Hattie Chung and Jenny Chen for insight on gene expression specificity analysis and Sam Myers for advice on proteomics. This work was funded by NIH Center for Excellence in Genomics Science, HHMI, the Klarman Cell Observatory, CZI Neurodegeneration Challenge Network, Open Philanthropy Project/Good Ventures, and work on “Brain-Derived Extracellular Vesicles for Analysis of Treatment Resistant Major Depressive Disorder” was supported by Wellcome Leap as part of the Multi-Channel Psych Program. AR was a Howard Hughes Medical Institute Investigator when this work was initiated.

## Contributions

DT conceptualized the study. DT, SW, TG, EJKK, WT, BB, CMB, and MWB performed experiments and analyzed data. DT, EJKK, SI, and GH performed computational analysis. HK, SAC, AR, GMC, and DRW supervised the study. DT wrote the manuscript with input from all authors.

## Materials and Methods

### Human biofluids

Both pooled and individual human plasma samples (collected in K2 EDTA tubes) were ordered from BioIVT. Pooled human CSF samples were ordered from BioIVT or Biochemed Services. Individual CSF samples were ordered from the Crimson Biomaterials Collection Core Facility at Brigham and Women’s Hospital. MMR Slurry, tetraspanin immuno-isolation method development and mass spectrometry was performed on pooled CSF from BioIVT. Candidate marker detection and neuron EV immuno-isolation was performed on pooled CSF from both BioIVT and Biochemed Services, as well as individual CSF from the Crimson Core. Human biofluid samples were thawed at room temperature and centrifuged for 10 minutes at 2000×*g*. Then, for all experiments other than MMR Slurry (where a filtration step is part of the protocol), samples were filtered through a 0.45-μm Corning Costar Spin-X centrifuge tube (MilliporeSigma) for 10 minutes at 2000×*g*.

### Size exclusion chromatography

Size exclusion chromatography (SEC) was performed as previously described using a custom-built stand (7, 16, 18). Sepharose CL-6B was washed three times in bulk with PBS before use. Econo-Pac Chromatography columns (Bio-Rad) were packed with Sepharose CL-6B and when the bed volume of dry resin reached 10 mL, a frit was inserted into the column above the resin. The resin was then washed in-column with 10 mL PBS before use. Sample (1 mL) was loaded onto the column once all PBS went through, and once sample went through, 2 mL PBS was added. Once all liquid went through (void volume), collection of 0.5 mL fractions began. For each fraction, 0.5 mL of PBS was added to top of column and then collected in 2 mL Eppendorf tube.

### EV isolation from plasma or CSF by MMR Slurry

Mixed mode chromatography resin slurry was prepared by taking Capto Core 700 resin (Cytiva) and washing 3 times with PBS in a 50 mL falcon tube. Resin was then resuspended in a volume equal to the resin volume of PBS to produce a 50% slurry. The protein concentration of the CSF or plasma SEC fractions 7-10 was then determined using Qubit Protein Assay kit (ThermoFisher Scientific) using 1 μL of sample. A volume of 50% Capto Core slurry corresponding to the protein content of the samples added. The ratio of slurry to total protein was varied, as specified (for example, for the CC0.6 condition, 1 μL of slurry was added per 0.6 μg of protein sample). Samples were mixed end over end for 45 minutes at room temperature, and then centrifuged at 800×*g* for 10 minutes. Finally, supernatant was transferred and centrifuged in Corning CoStar X 0.45 μm filters (MilliporeSigma) at 2000×*g* for 10 minutes to separate the CSF from the Capto Core beads. Samples were concentrated at 4°C for 90 minutes using Amicon Ultra-2 centrifugal 10 kD filter (MilliporeSigma) at 3000×*g*.

### Mass spectrometry

Mass spectrometry for cell culture EVs was performed at the Broad Institute Proteomics Platform. iPS and iNGN EVs were lysed in RIPA buffer (ThermoFisher Scientific) and two sets of triplicate technical replicates were run on NuPage 4-12% Bis-Tris SDS Page (ThermoFisher Scientific) gels in 10 mM DTT/1x loading buffer. Gel was run at 130V with 130mA for 50 min followed by Coomassie stain. Gel bands were excised into 6 bands whereby the intense albumin band was kept separate from the remaining bands to improve sensitivity. In-gel digestion was performed on all 72 bands via rehydration of gel pieces with 25 mM ammonium bicarbonate (ABC) followed by reduction with 10 mM DTT (56°C for 1 hour) and alkylation with 55mM iodoacetamide (45 min at room temperature in the dark). Gel pieces were then washed, dehydrated with 25 mM ABC/50% acetonitrile, and rehydrated with Trypsin at 37°C overnight. Peptides were then incubated at RT for 30 min, extracted with 50% acetonitrile/5% formic acid, and combined with the supernatant fraction. Samples were then dried via vacuum centrifugation and then desalted via stage-tip desalting. Samples were reconstituted in 10μL of 5% acetic acid/3% acetonitrile with 4 μL injected. Each sample was run on a NanoLC (Easy LC; ThermoFisher Scientific) coupled with a Q Exactive Plus (ThermoFisher) mass spectrometer with the following parameters: 110 min method with effective gradient of 0 to 30%B acetonitrile at 0.35%/min followed by 30-60%B at 3.3%/min at 200nL/min.

Mass spectrometry for EVs isolated by MMR Slurry was performed at the Harvard Center for Proteomics. EV protein was ethanol precipitated and submitted as a protein pellet. Samples were precipitated by adding 9 volumes of cold 100% ethanol to 1 volume of EVs, vortexed for one minute, and kept at −20°C for 30 minutes. Samples were then centrifuged at 16,000×*g* for 15 minutes at 4°C. Supernatants were removed and pellets were left to air dry for 10 minutes. Protein was resuspended in TEAB and digested using Filter Aided Sample Preparation (FASP) (52) on a 10 kDa filter (Pall). After drying samples down in a SpeedVac (Eppendorf), digested protein was resuspended in 0.1% formic acid and submitted for LC-MS/MS on a Orbitrap Fusion Lumos Tribrid (ThermoFisher Scientific) equipped with a 3000 Ultima Dual nanoHPLC pump (ThermoFisher Scientific). Peptides were separated onto gradient from 5–27% ACN in 0.1% formic acid over 90 min at 200 nl min^−1^, instrument was operated in data-dependent acquisition (DDA) mode.

### Mass spectrometry data analysis

Data analysis for cell culture EVs was done using the Spectrum Mill MS Proteomics Workbench software package v 4.2 beta (Agilent Technologies). All extracted spectra were searched against a UniProt database containing human reference proteome sequences. Database matches were validated at the peptide and protein level in a two-step process with identification FDR estimated by target-decoy-based searches using reversed sequences.

For plasma and CSF EV proteomes data analysis, raw data from several samples of EVs isolated from pooled plasma or CSF (all from BioIVT) were combined and submitted for analysis in Proteome Discoverer 2.4 (ThermoFisher Scientific). Assignment of MS/MS spectra was performed using the Sequest HT algorithm by searching the data against a protein sequence database including all entries from the Human Uniprot (2019 release) database (53) and other known contaminants such as human keratins and common lab contaminants. A MS2 spectra assignment false discovery rate (FDR) of 1% on protein level was achieved by applying the target-decoy database search and filtering was performed using Percolator (54).

### Cell type-specific expression by bulk RNA-Seq

For Tau calculations, only genes that had Log2(TPM+1)>0.3 for at least one cell type or organ were selected. Organ-specific Tau was calculated (24) using GTEx organ-specific RNA-seq data (23), after removing pituitary gland, tibial nerve, and testis samples. Since GTEx data contains several regions or tissues for each organ, all regions or tissues for a specific organ were averaged, resulting in one organ-level measurement. Brain cell type-specific Tau was calculated using Brain RNA-Seq expression data of the five main cell types of the brain: neuron, astrocytes, oligodendrocytes, microglia, and endothelial cells (22). To determine genes which were specific to neurons among brain-cell types, and specific to the brain within all organs, we selected genes that were: expressed at higher levels in neurons relative to other brain cell-types in Brain RNA-seq data, expressed at higher levels in brain relative to other organs in GTEx data, and had a Tau score of 0.7 in both.

### Cell type-specific expression by scRNA-Seq

SCimilarity’s Interpreter class (26) was used to identify genes that distinguish two groups of cells. This was used to find marker genes for astrocytes, oligodendrocytes, microglia, and neurons. To find genes that distinguished each of these cell types from all other cell types, 1000 cells were randomly sampled from SCimilarity’s 399 reference studies that were predicted as cell type of interest (anchor cells) and 1000 cells that were not of the target cell type nor any descendant cell types based on Cell Ontology relationships (negative cells). To find genes that were high in our anchor population and low in all other cell types, integrated gradients were applied to associate input features (*i.e.* genes) with SCimilarity’s low dimensional embeddings. Genes with the highest attributions scores were most important in distinguishing between anchor and negative cell groups. Genes that SCimilarity learns to be important for cells’ embeddings receive higher integrated gradient scores, while genes that the model is less sensitive to receive scores near zero.

### Selection of candidate EV cell type-specific markers for neurons

Uniprot accession IDs for all human proteins were filtered for those annotated to contain a transmembrane domain (55). The final candidate markers were selected as genes that (**1**) had an annotated transmembrane domain; (**2**) were present in the EV CSF or plasma proteomics data; and (**3**) had a Tau score of 0.7 or greater for both neuron-specific expression in Brain RNA-Seq and brain-specific expression in GTEx and/or had an SCimilarity Attribution Score of 0.001 or greater for neurons.

### Cell culture

K562 cells (from ATCC) were grown in Gibco IMDM with Glutamax (ThermoFisher Scientific) supplemented with Gibco Heat-Inactivated Fetal Bovine Serum (ThermoFisher Scientific) and Gibco Penicillin Streptomycin (Thermo Fisher Scientific). Previously described iNGN cells (40) were grown in mTeSR1 media (STEMCELL Technologies) on Matrigel (Corning) coated plates. Doxycycline (Sigma Aldrich) was diluted in PBS and added to mTeSR1 at a final concentration of 0.5 μg/mL to initiate differentiation. On Day 4 after Dox addition, media was switched to Gibco DMEM with Glutamax (ThermoFisher Scientific) supplemented with B27 Serum-Free Supplement (ThermoFisher Scientific) and Gibco Penicillin Streptomycin (ThermoFisher Scientific). A detailed protocol is available: dx.doi.org/10.17504/protocols.io.bn63mhgn

### EV isolation from cell culture media

For K562 EV isolation, K562 cells were switched to EV-depleted media (obtained by ultracentrifugation of media for 16 hours at 120,000xg and subsequent filtration through Corning 0.22 μm filter). EVs from iNGN neurons were collected on Day 6 or Day 7 after Dox addition. EV isolation from cell culture was performed by differential ultracentrifugation as previously described (41). Briefly, cell culture media (240 mL per isolation) was centrifuged at 300×*g* for 10 minutes and supernatant was centrifuged again at 2000×*g* for 10 minutes. Supernatant was centrifuged at 16,500×*g* at 4°C for 20 minutes and filtered through 0.22μm Steriflip filter (MilliporeSigma). Samples were then ultracentrifuged at 120,000×*g* at 4°C for 70 minutes, washed with PBS and ultracentrifuged again. The pellet was then resuspended in PBS. A detailed protocol is available: dx.doi.org/10.17504/protocols.io.bnr3md8n

### Total protein staining

Protein samples were denatured in LDS (ThermoFisher Scientific) for 10 minutes at 70°C before loading on a Bolt Bis-Tris Plus 4 to 12% gel (ThermoFisher Scientific) for total protein staining or Western blotting. Samples were run at 150V for 60 minutes. Coomassie Blue total protein staining was performed on gels using Acqua Stain (Bulldog Bio). The gel was incubated in the stain overnight, washed in deionized water, and then imaged with a Gel Doc EZ Imager (BioRad).

### Western blotting

Western blotting of EVs was performed as previously described (46). Briefly, the iBlot2 Dry Blotting System (ThermoFisher Scientific) was used for transfer at 20V for three to seven minutes, depending on the size of the protein marker. The following primary antibodies were used for Western blot at the corresponding dilutions: M38 for CD81 (Thermo Fisher Scientific) at 1:666, H5C6 for CD63 (BD) at 1:1000, CD9 (Millipore) at 1:1000, EPR18998 for L1CAM (Abcam) at 1:500, ab47441 for GJA1 at 1:500, F-10 for albumin (Santa Cruz) at 1:1000, E7K3G (Cell Signaling Technology) at 1:5000 or AF5269 (R&D Systems) at 1:1000 for NRXN3. Blots were incubated on a shaker in milk (5% weight by volume) dissolved in PBS-T solution (PBS with 0.1% Tween) containing the primary antibodies overnight at 4°C. The next day, blots were washed thrice with PBS-T, incubated with TrueBlot HRP (Rockland) or cross-adsorbed HRP (Bethyl) secondary antibody at a concentration of 1:2000 in milk buffer for 2 hours, and then washed thrice again. Blots were developed with WesternBright ECL-spray (Advansta) and imaged on a Sapphire Biomolecular Imager (Azure Biosystems). A detailed protocol is available: dx.doi.org/10.17504/protocols.io.bnrtmd6n

### EV immuno-isolation from cell culture EVs

Isolation buffer was prepared by adding BSA to 7.4 pH PBS to final concentration of 1mg/mL and filtered through a 0.22 μm Steriflip filter (Millipore). 500 μL (2x10^8^) Dynabeads Goat Anti-Mouse IgG beads (ThermoFisher Scientific) were put into 2mL and placed on a magnetic rack. Supernatant was removed and replaced with 250 μL of isolation buffer off the magnet. 10 μg primary antibody was coupled to beads overnight at 4°C with end over end rotation. The following antibodies were used for immuno-isolation: 5G3 (BD) for L1CAM, 1C51 for mCherry (Abcam), 9F9.F9 for GFP (Abcam), 1.3.3.22 (ThermoFisher Scientific) for CD81, and H5C6 (BD) for CD63. On the next day, beads were washed twice with 1 mL of isolation buffer each time. EVs were then added (usually one pellet was in 150 μL) and isolation buffer was added to bring the volume to 0.5 mL. Immuno-isolation was performed on rotating rack either at 4°C for 24 hours for L1CAM or for 1 hour at 37°C for CD81 or CD63.

### EV immuno-isolation from human biofluids

EV immuno-isolation from CSF or plasma was performed similarly to EV immuno-isolation from cell culture EVs (described above), with a few minor modifications. PBS pH 7.4 was used without addition of BSA as the isolation buffer. 250 μL (1x10^8^) Dynabeads Goat Anti-Mouse IgG beads (ThermoFisher Scientific) or 50 μL (1.5 mg) of Dynabeads Protein A (ThermoFisher Scientific) were dispensed into a 2 mL tube and placed a magnetic rack. Supernatant was removed, beads were washed with 1 mL isolation buffer, and then brought up to final incubation volume of 0.5 mL with PBS pH 7.4 and 10 μg primary antibody. Antibody was coupled to beads rotating end over end overnight at 4°C. The following antibodies were used for immuno-isolation: mouse monoclonal CD81 (clone 1.3.3.22, ThermoFisher Scientific), CD63 (clone H5C6, BD Biosciences), CD9 (clone CBL162, Millipore), GFP (clone 1GFP63, BioLegend or clone 9F9.F9, Rockland) and mCherry (clone EPR20579, Abcam). The next day, beads were washed twice with 1 mL isolation buffer. The CSF, plasma, or SEC (0.5 mL input) immuno-isolation was performed on a rotating rack for 1 hour at 4°C for CD9, CD63 and CD81. The following antibodies were used for immuno-isolation of NRXN3 and controls: sheep polyclonal NRXN3 (cat. AF5269, R&D Systems), GFP (4745-1051, Rockland), and Isotype Control (AB37385, Abcam). To couple sheep antibody to beads, 250 μL (1x10^8^) Dynabeads M-450 Epoxy (14011, ThermoFisher Scientific) were incubated overnight at 4°C with the antibody. Antibody-coupled beads were washed twice with 1 mL of isolation buffer and mixed with 0.5 mL of biofluid or SEC fractions 7-10 on a rotating rack for 24 hours at 4°C. Post immuno-isolation, biofluid was collected from the beads, along with two isolation buffer washes of 0.75 mL each. Collected “flow through” was diluted and measured by Simoa.

### Simoa

Simoa for CD9, CD63, CD81, and albumin was performed as previously described (7, 16, 18). A detailed protocol is available: https://www.protocols.io/view/simoa-extracellular-vesicle-assays-81wgb7pw1vpk/v1. New assays were created by cross-testing all commercially available antibodies for NRXN3, SLC32A1, and SLC12A5 (**Table S7**). For each marker, 3 pairs of antibodies with the highest signal to noise ratio were selected and used to test spike and recovery, dilution linearity, and signal against cell culture neuron EV SEC fractions. For spike and recovery, a dilution factor in the linear range during dilution linearity was chosen to be the dilution factor and protein standard was spiked into plasma or CSF. Recoveries of spike within 70-130% were considered successful. The pair which performed best on all three tests was optimized for Simoa. For NRXN3, AF5269 (R&D Systems) was used as capture and 47714 (ThermoFisher Scientific) was used as detector. Purified human recombinant NRXN3 TP323448 (Origene) was used as standard. For SLC12A5, MAB8369 (R&D Systems) was used as capture and 821701 (BioLegend) was used as detector. TP323680 (Origene) was used as standard. For SLC32A1, NB110-55238 (Novus Biologicals) was used as capture, and mab6847 was used as detector (Novus Biologicals). TP762191 (Origene) was used as standard. Homebrew detector/ Sample diluent (Quanterix) was used as diluent in all neuron marker assays. Duplicates were measured for each sample on the HD-X Analyzer (Quanterix) and average enzyme on bead (AEB) was calculated by HD-X software (Quanterix).

### Density gradient centrifugation

Density gradient centrifugation (DGC) was performed as previously described (18). Four layers of OptiPrep (iodixanol) were prepared and arranged in 13.2 mL propylene tubes sequentially from bottom to top: 3 mL 40%, 3mL 20%, 3mL 10%, 2mL 5%. OptiPrep (MilliporeSigma) was diluted in a solution of 0.25M sucrose (MilliporeSigma) and pH 7.4 Tris–EDTA (MilliporeSigma). Sample was loaded on top of the gradient and centrifuged at 100,000 RCF in an SW 41 Ti swinging bucket rotor for 18 hours at 4°C using a Beckman Coulter Optima XPN-80 ultracentrifuge. Fractions were then removed from the top 1 mL at a time.

## Data availability

Mass spectrometry data is available in Supplementary Information Tables S1 and S8. Raw data will be uploaded to ProteomeXchange upon publication.

## Code availability

Python scripts for the computational marker pipeline are available: https://github.com/Wyss/extracellular-vesicle-markers.

