## Supplemental Information for "Identification of markers for the isolation of neuron-specific extracellular vesicles"

### Supplementary Information

Supplementary Table S1: List of proteins detected by mass spectrometry in EVs isolated by MMR Slurry from CSF and plasma

Supplementary Table S2: List of all proteins with transmembrane domains

Supplementary Table S3: List of proteins that are brain and cell type-specific by bulk RNA-Seq and whether they have transmembrane domains

Supplementary Table S4: List of sorted proteins by SCimilarity Attribution Scores to quantify specificity from scRNA-Seq data for neurons, astrocytes, oligodendrocytes, and microglia and whether they have transmembrane domains

Supplementary Table S5: List of proteins that are cell type specific by both bulk RNA-Seq and scRNA-Seq

Supplementary Table S6: List of candidate EV markers that are transmembrane and specific to brain and neurons, astrocytes, oligodendrocytes, or microglia by bulk RNA-Seq with other attributes

Supplementary Table S7: List of all antibodies tried for development of Simoa assays for candidate neuron markers

Supplementary Table S8: Mass spectrometry data of proteins found on EVs isolated from cell culture media of iPS cells and iNGN neurons

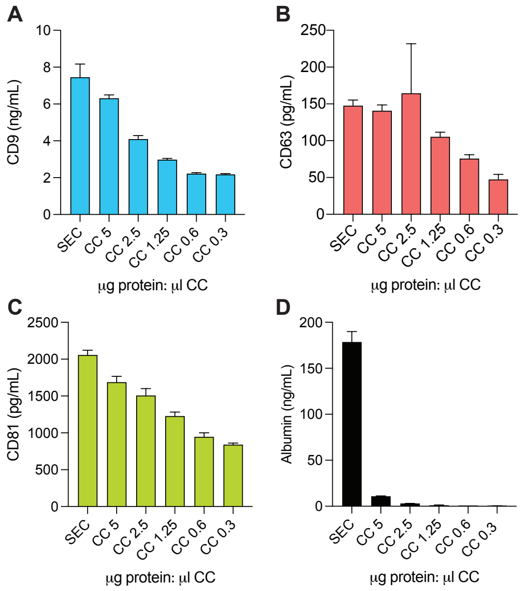


**Figure S1: MMR Slurry isolation significantly reduces concentration of albumin relative to tetraspanins**

Simoa measurement of **A.** CD9 **B.** CD63 **C.** CD81 **D.** and albumin measured after isolation of EVs from 0.5 mL plasma by SEC or SEC followed by MMR Slurry with increasing amounts of Capto Core beads. Two replicates were performed for each condition and Simoa measurements were taken in duplicate for each replicate. Error bars represent standard deviation from the mean.

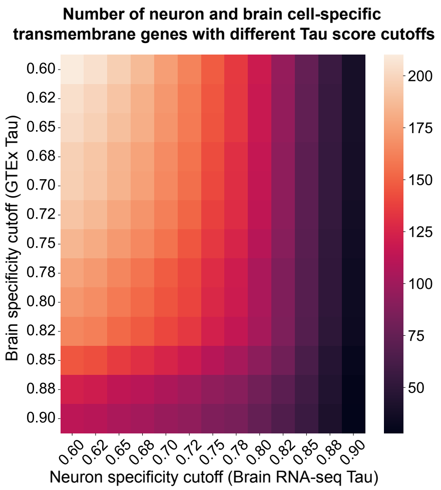


**Figure S2: Heatmap of Tau Cutoffs for bulk RNA-Seq**

Number of transmembrane proteins when different cutoffs are used of Tau specificity score for neurons within the brain (using Brain RNA-Seq) and the brain relative to other organs of the body (using GTEx).

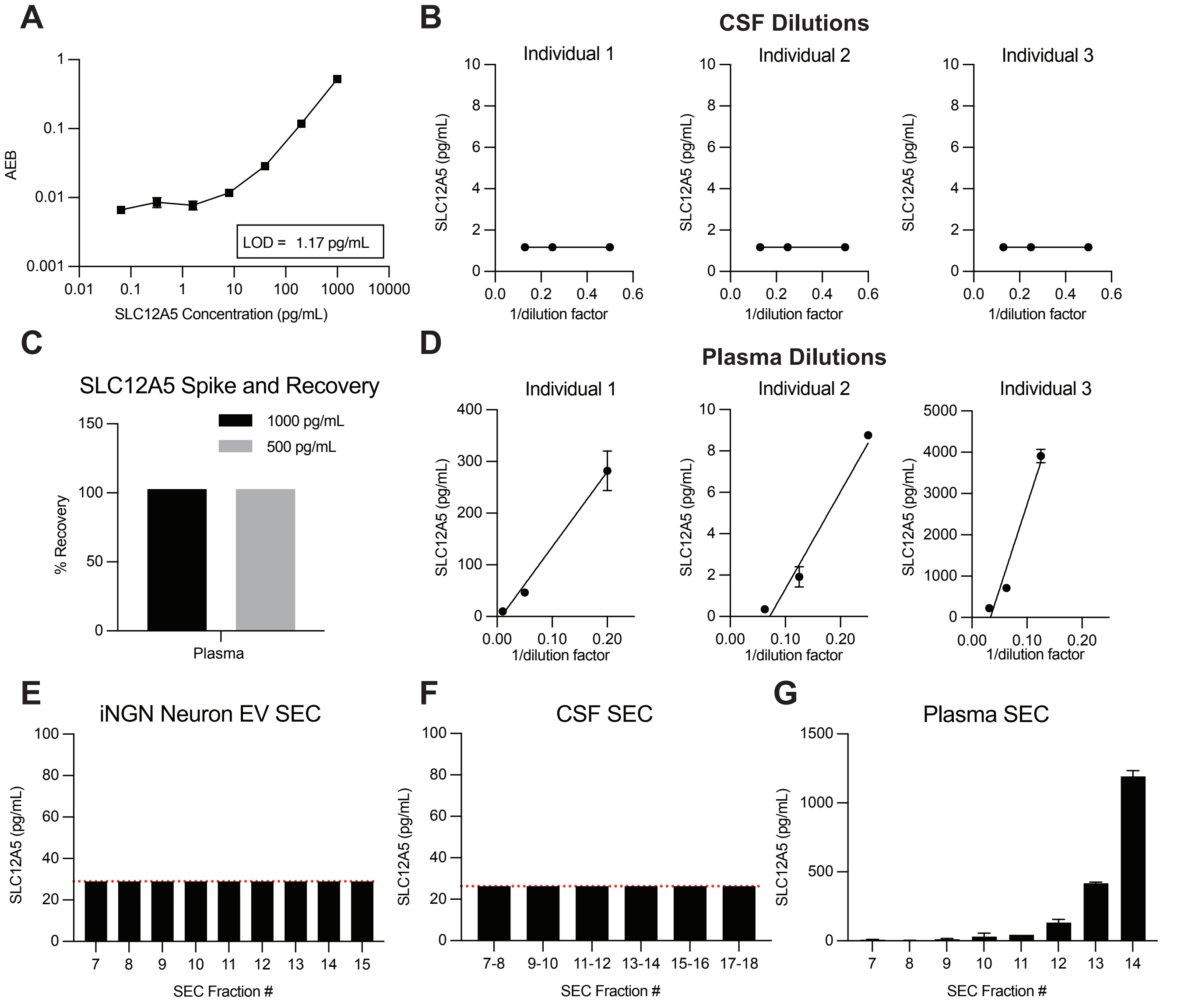


**Figure S3: Development of Simoa assay for SLC12A5**

**A.** Simoa measurement of calibration curve with purified SLC12A5 protein. Limit of detection (LOD) was determined to be 1.17 pg/mL. **B.** CSF dilutions from three individuals (all below LOD). **C.** High (500pg) and low (100pg) spike of purified SLC12A5 protein into plasma measuring percent recovery. **D.** Plasma dilutions from three individuals. **E.** Simoa Measurement of SEC fractions from EVs enriched from the cell culture media of iNGN neurons by differential ultracentrifugation (below LOD). **F.** Simoa measurement of SEC fractions from CSF (below LOD). **G.** Simoa measurement of SEC fractions from plasma. Simoa measurements were all performed in duplicate. Error bars represent standard deviation from the mean.

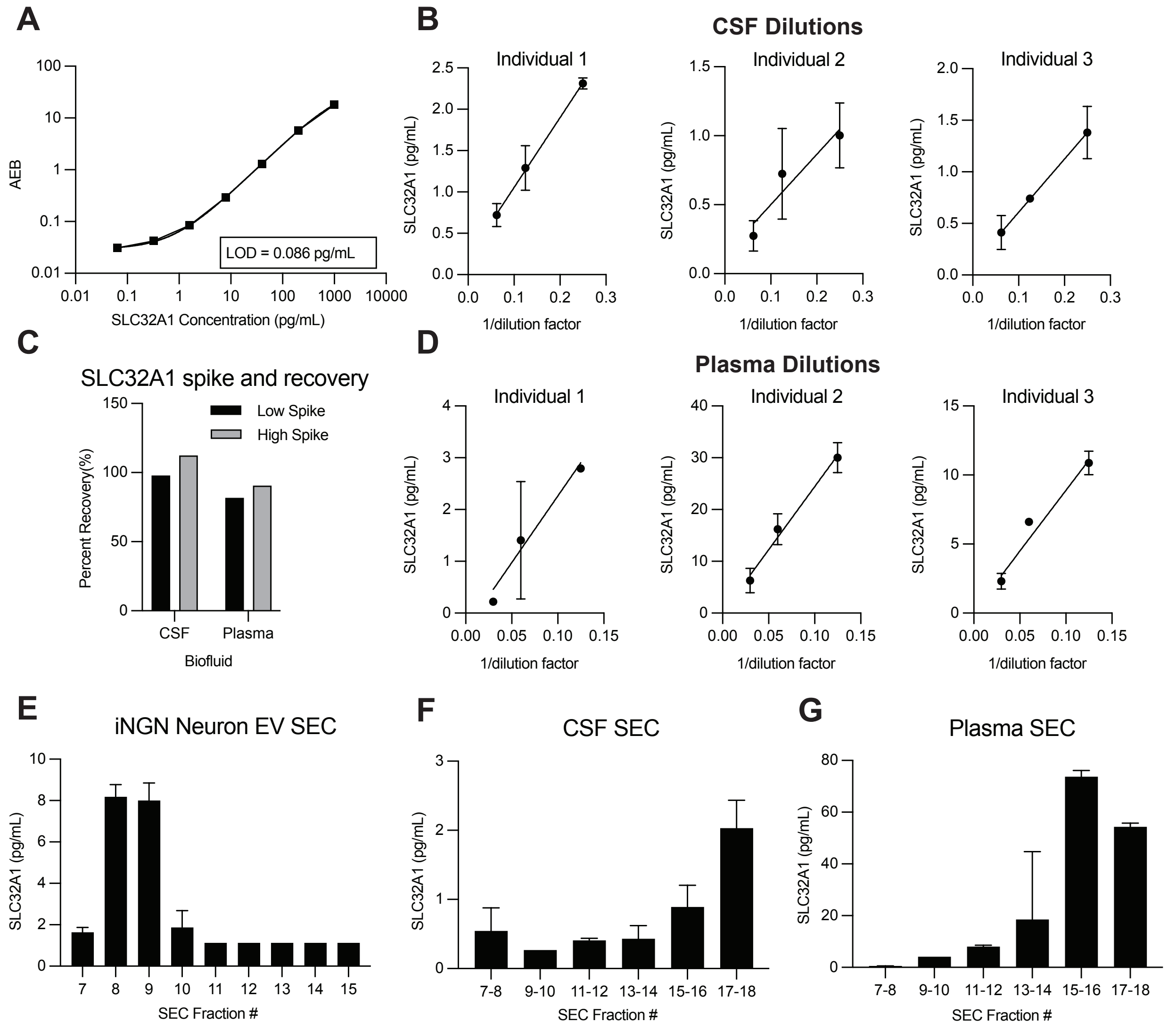


**Figure S4: Development of Simoa assay for SLC32A1**

**A.** Simoa measurement calibration curve with purified SLC32A1 protein. Limit of detection (LOD) was determined to be 0.086 pg/mL. **B.** CSF dilutions from three individuals (all below LOD). **C.** High (500pg) and low (50pg) spike of purified SLC32A1 protein into plasma measuring percent recovery. **D.** Plasma dilutions from three individuals. **E.** Simoa measurement of SEC fractions from EVs enriched from the cell culture media of iNGN neurons by differential ultracentrifugation. **F.** Simoa measurement of SEC fractions from plasma. **G.** Simoa measurement of SEC fractions from CSF. Simoa measurements were all performed in duplicate. Error bars represent standard deviation from the mean.

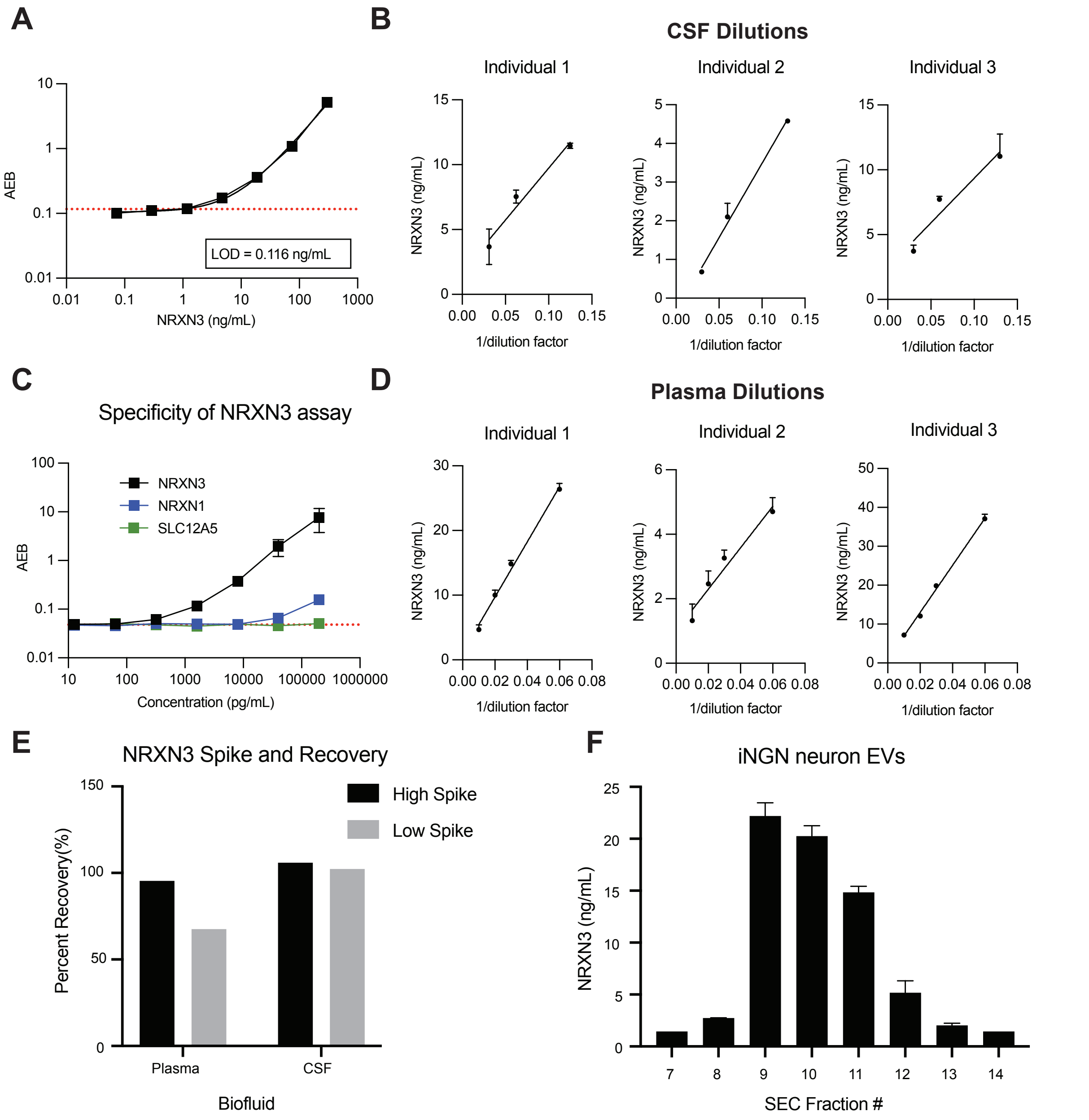


**Figure S5: Development of Simoa assay for NRXN3**

Simoa measurement of calibration curve with purified NRXN3 protein. Limit of detection (LOD) was determined to be 0.116 ng/mL. **B.** CSF dilutions from three individuals. **C.** Calibration curve with NRXN3 (black), NRXN1 (blue) or SLC12A5 (green) purified protein standard to assess the specificity of the NRXN3 Simoa assay for NRXN3. **D.** Plasma dilutions from three individuals. **E.** High (500ng) and low (50ng) spike of purified protein NRXN3 into plasma and CSF, measuring percent recovery relative to spike into diluent. **F.** Simoa measurement of SEC fractions from EVs enriched from the cell culture media of iNGN neurons by differential ultracentrifugation. Simoa measurements were all performed in duplicate. Error bars represent standard deviation from the mean.

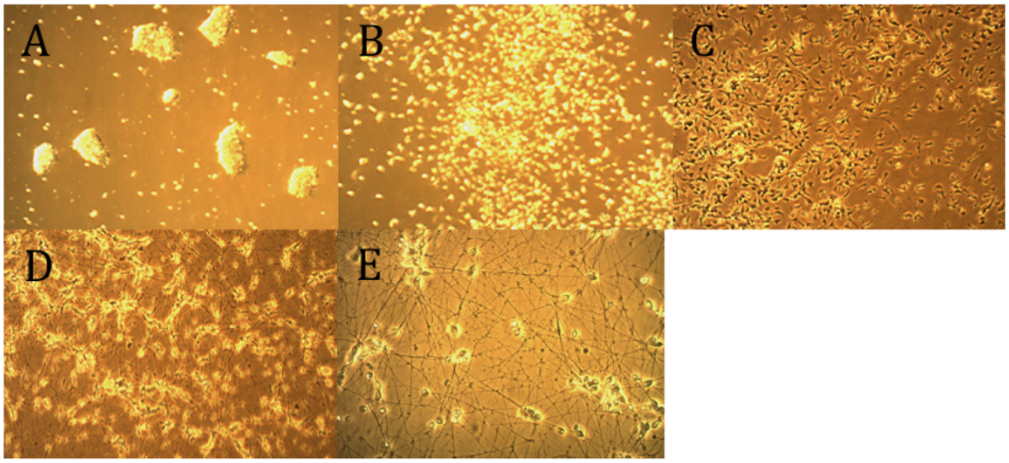


**Figure S6: Differentiation of iNGN cells from iPS to neurons upon doxycycline addition**

**A.** iPS iNGN cells before doxycycline addition (40x magnification). **B.** iPS iNGN cells 1 day after doxycycline addition (40x magnification). **C.** iPS iNGN cells 2 days after doxycycline addition (40x magnification). **D.** iPS iNGN cells 4 days after doxycycline addition (40x magnification) **E.** iNGN cells 5 days after doxycycline addition (200x magnification).

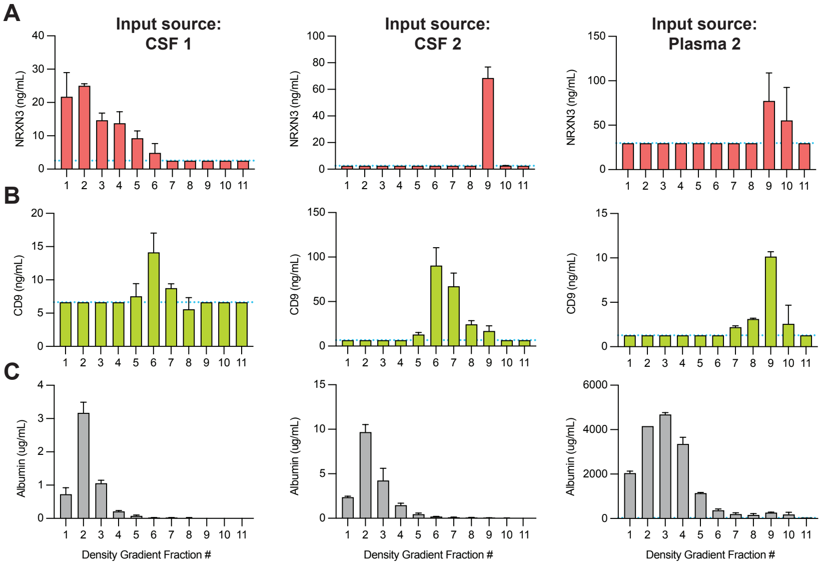


**Figure S7: Simoa of NRXN3 in pooled CSF and plasma fractionated by density gradient centrifugation**

Density gradient centrifugation (DGC) fractions for two different pooled CSF samples (first and second row) and a pooled plasma sample (third row, different pool than that shown in Figure 3C) were analyzed by Simoa for: **A.** NRXN3 **B.** CD9 and **C.** albumin. Simoa measurements were all performed in duplicate. Error bars represent standard deviation from the mean.

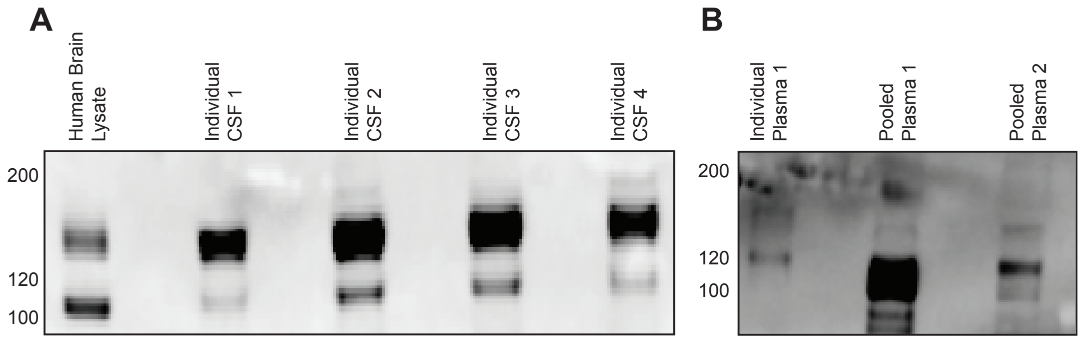


**Figure S8: Western blot of NRXN3 MMR Slurry**

**A.** Western blot of NRXN3 in four individual CSF samples after MMR slurry purification with 2.5 µg protein: µL resin. Human brain lysate is loaded as a positive control. **B.** Western blot of NRXN3 in individual and pooled plasma samples after MMR slurry purification with 0.6 µg protein: µL resin.

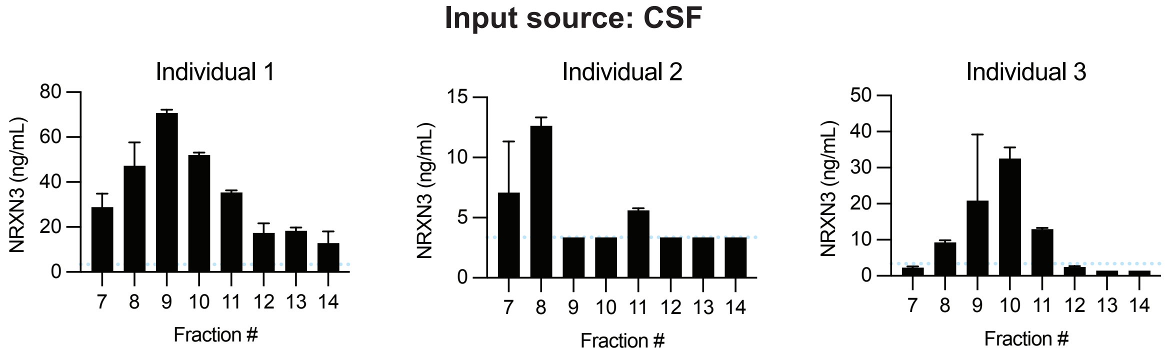


**Figure S9: NRXN3 peaks in EV fractions in SEC across individual CSF**

Simoa measurement of NRXN3 in SEC fractions 7-14 from three individual CSF samples. Limit of detection shown in blue. Simoa measurements were all performed in duplicate. Error bars represent standard deviation from the mean.

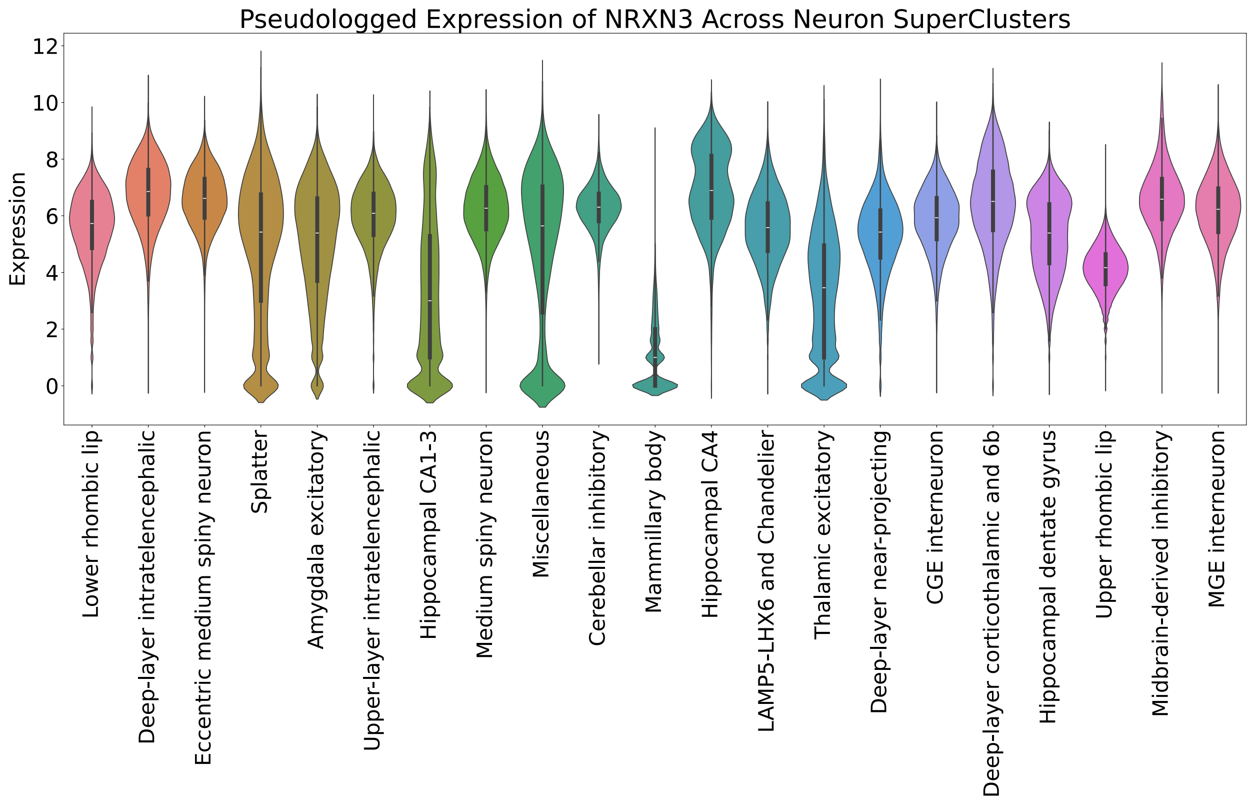


**Figure S10: Expression of NRXN3 across neuronal subtypes in the human brain**

NRXN3 expression in single cells in the human brain across neuron cell subtypes annotated in Siletti *et al*, 2023. Expression is plotted as log2(UMI Counts+1).

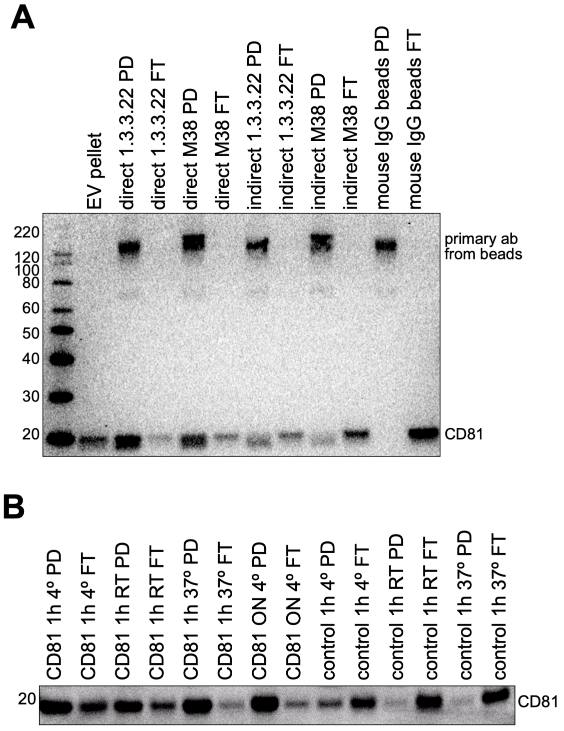


**Figure S11: Optimization of CD81 immuno-isolation parameters using EVs from K562 cells**

**A.** Western blot of CD81 after EVs isolated by differential ultracentrifugation were immuno-isolated using CD81 antibody. Direct (primary antibody conjugated to beads and then added to sample) vs. indirect (primary antibody is added to sample first and then added to beads) pulldown methods are compared for CD81 immuno-isolation using two different antibodies (clones M38 and 1.3.3.22). Note: band at 150 kDa is primary antibody coming off the beads. **B.** Western blot against CD81 after EVs were incubated with beads for either 1 hour or overnight (ON), at different temperatures (4°C, room temperature, or 37°C) using beads conjugated to either CD81 or a control non-specific antibody (against mCherry).

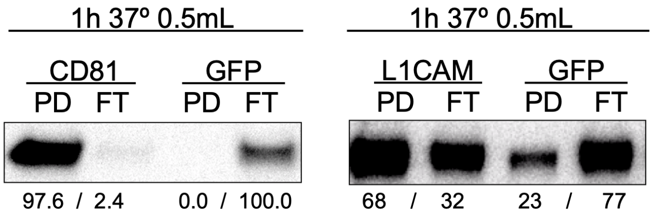


**Figure S12: Optimal immuno-isolation conditions vary for different antibodies**

Immuno-isolation for 1 hour at 37°C in 0.5 mL volume was performed on EVs isolated by differential ultracentrifugation from the conditioned media of K562 cells using CD81 (or control GFP antibody) and neuron EVs using L1CAM (or control GFP antibody) and corresponding Western blot was performed against CD81 or L1CAM. Intensities of bands on Western blot were quantified using ImageJ as percentage of band relative to sum of pulldown (PD) + flow-through (FT) for that condition.

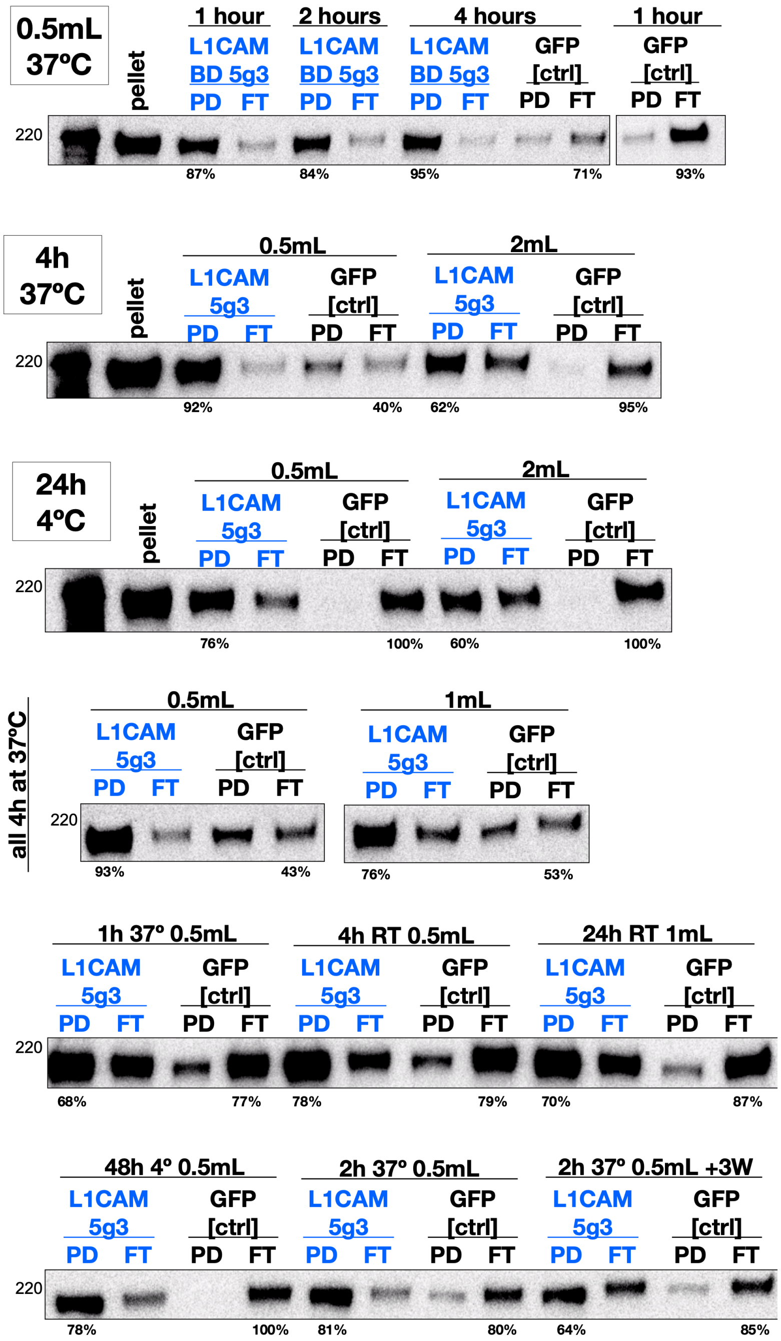


**Figure S13: Optimization of conditions for iNGN neuron EV immuno-isolation**

Western blot against L1CAM after iNGN neuron EVs were incubated with antibody conjugated beads (against L1CAM or GFP) for different times (1, 2, 4, or 24 hours), in different volumes (0.5 mL or 2 mL) at different temperatures (4°C or 37°C) and with different numbers of wash steps (+3W). Intensity of bands on Western blot was quantified using ImageJ as percentage of band relative to sum of pulldown (PD) + flow-through (FT) for that condition.

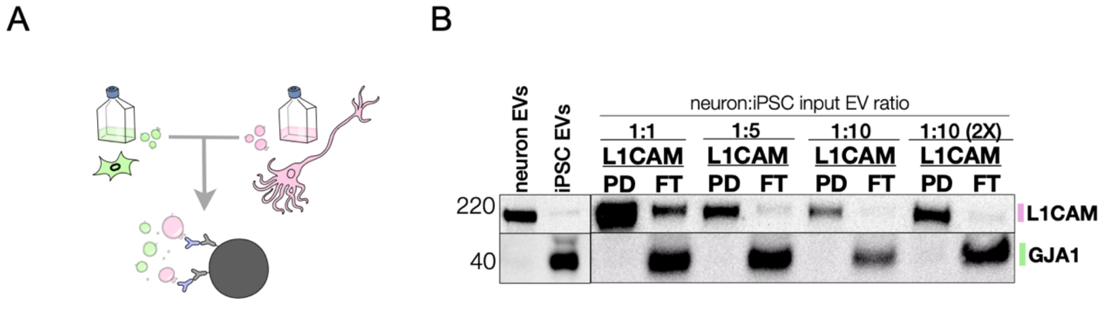


**Figure S14: iNGN neuron EV immuno-isolation from iPS/iNGN EV mixture**

**A.** Mixing EVs isolated by differential ultracentrifugation from the conditioned media of iPS cells and iPS-derived iNGN neurons allows for assessment of pulldown efficiency and purity in a more complex mixture model. **B.** Western blot of L1CAM and GJA1 after L1CAM immuno-isolation demonstrates that the immuno-isolation efficiently and specifically isolates L1CAM+ neuron-derived EVs from pooled iNGN neuron EVs and iPS EVs in varying ratios (normalized relative to cell number). An iPS specific marker, GJA1, is analyzed to ensure iPS EVs are not non-specifically bound to L1CAM beads. L1CAM was found on iNGN neuron EVs by mass spectrometry and we have previously found L1CAM to be present on iNGN EV, unlike on plasma and CSF EVs, by SEC (Norman *et al*, 2021).

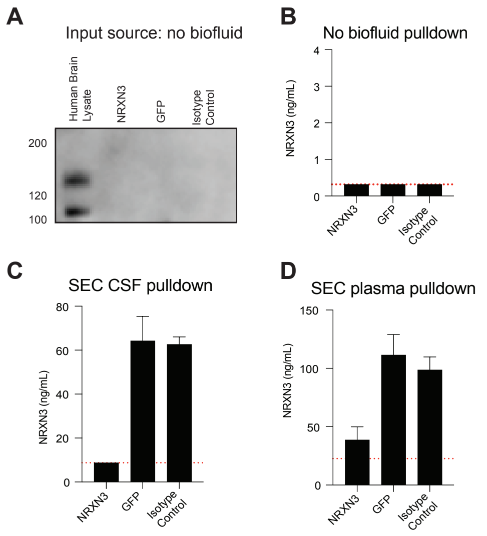


**Figure S15: Immuno-isolation of NRXN3 from human CSF and plasma after SEC**

**A.** Western blot of NRXN3 after incubation of beads without biofluid (as a negative control). **B.** Simoa measurement of NRXN3 after incubation of immunoprecipitation beads without biofluid (as a negative control). **C.** Simoa measurement of NRXN3 concentration after incubation of NRXN3 beads or control GFP/ isotype control beads with CSF fractions 7-10 collected from SEC. **D.** Simoa measurement of NRXN3 from plasma fractions 7-10 collected after SEC. Limit of detection for the assay shown in red. Simoa measurements were all performed in duplicate. Error bars represent standard deviation from the mean.
